# Predicting MHC-I ligands across alleles and species: How far can we go?

**DOI:** 10.1101/2024.05.08.593183

**Authors:** Daniel M. Tadros, Julien Racle, David Gfeller

## Abstract

CD8^+^ T-cell activation is initiated by the recognition of epitopes presented on class I major histocompatibility complex (MHC-I) molecules. Identifying such epitopes is useful for molecular understanding of cellular immune responses and can guide the development of personalized vaccines for various diseases including cancer. Here, we capitalize on high-quality MHC-I peptidomics data available from different species and an expanded architecture of our MHC-I ligand predictor (MixMHCpred) to carefully explore how much predictions can be extrapolated to MHC-I alleles without known ligands. Our results reveal high prediction accuracy for most MHC-I alleles in human and in laboratory mouse strains, but significantly lower accuracy in other species. Our work further outlines some of the molecular determinants of MHC-I ligand predictions accuracy across alleles and species. Robust benchmarking on external data shows that the pan-allele version of MixMHCpred (v3.0) outperforms other state-of-the-art MHC-I ligand predictors and can be used for CD8^+^ T-cell epitope predictions.

## Introduction

CD8^+^ T cells play a key role in eliminating infected or malignant cells. To perform this task, CD8^+^ T cells recognize small peptides displayed on class I major histocompatibility complex (MHC-I) molecules on the surface of the targeted cells. MHC-I ligands are considered as promising therapeutic targets and have been used in pre-clinical and clinical studies. For instance, in cancer immunotherapy, MHC-I ligands have been used as personalized vaccines to boost the immune system to recognize neo-antigens (Carreno et al., 2015; Leidner et al., 2022; Migliorini et al., 2019; Ott et al., 2017). Additionally, viral peptides presented on MHC-I molecules have been utilized in vaccines against infectious diseases to elicit strong T-cell recognition (Heitmann et al., 2022).

MHC-I molecules bind short peptides (roughly 8–14 amino acids) with a general preference for 9-mers (Gfeller et al., 2018a; Lundegaard et al., 2008; Nielsen & Andreatta, 2016; Trolle et al., 2016). The binding is typically determined by primary anchor residues at the second and last positions of the peptides. Several alleles display additional anchor residues at other positions (Gfeller et al., 2018a; Gfeller & Bassani-Sternberg, 2018). In human, MHC-I molecules are encoded by three commonly expressed genes (HLA-A, HLA-B, HLA-C) along with a few other genes (e.g., HLA-E, HLA-F, HLA-G). MHC-I genes exhibit a very high level of polymorphism, with thousands of distinct alleles (Barker et al., 2023). MHC-I genes in various species are known to evolve rapidly and are not strongly conserved, even among relatively closely related species (Bernatchez & Landry, 2003; Piertney & Oliver, 2006; Sommer, 2005). Different MHC-I alleles have different peptide-binding specificities, which include differences in binding motifs and peptide length distributions (Gfeller et al., 2018b; Tadros et al., 2023; Trolle et al., 2016). This results in distinct repertoires of MHC-I ligands in different individuals and different species (Gfeller, Liu, et al., 2023; Neefjes et al., 2011; Thibault & Perreault, 2022; Vyas et al., 2008). Over the last decade, mass spectrometry-based MHC-I peptidomics has emerged as the leading source of information about MHC-I binding specificities. These data have enabled researchers to compute binding motifs and peptide length distributions for hundreds of MHC-I alleles (Abelin et al., 2017; Bassani-Sternberg et al., 2017a; Pyke et al., 2021; Sarkizova et al., 2020).

Many in silico prediction tools for MHC-I ligands have been developed to narrow down the list of potential epitopes (Bulik-Sullivan et al., 2019; Gfeller, Schmidt, et al., 2023; O’Donnell et al., 2020; Pyke et al., 2021; Reynisson et al., 2020; Sarkizova et al., 2020). These tools are mainly trained on MHC-I peptidomics data and such data are currently available for a bit more than hundred alleles (Gfeller, Schmidt, et al., 2023; Nielsen et al., 2018; Sarkizova et al., 2020; Tadros et al., 2023; Vita et al., 2019). These include all common alleles in human but only a few alleles from other species. Two main classes of MHC-I ligand predictors can be distinguished: allele-specific or pan-allele predictors. Allele-specific predictors, such as NetMHC (Andreatta & Nielsen, 2016) or MixMHCpred2.2 (Gfeller, Schmidt, et al., 2023), train a separate model for each allele with known ligands. These tools are therefore restricted to the set of MHC-I alleles with available data. Pan-allele predictors, like NetMHCpan4.1 (Reynisson et al., 2020), MHCflurry2.0 (O’Donnell et al., 2020), ACME (Hu et al., 2019), MATHLA (Ye et al., 2021), DeepLigand (Zeng & Gifford, 2019), HLAthena (Sarkizova et al., 2020), BigMHC (Albert et al., 2023) or SHERPA (Pyke et al., 2021) can make predictions for a broader range of alleles. These predictors leverage shared properties of MHC-I molecules across different alleles and correlations between MHC-I binding site residues and binding specificities to make predictions even when experimental ligands are not available for a given allele. Pan-allele MHC-I ligand predictors have been successfully used in multiple studies in human (Hu et al., 2019; O’Donnell et al., 2020; Reynisson et al., 2020; Ye et al., 2021; Zeng & Gifford, 2019). However, considering the rapid evolution of MHC-I genes and alleles across species, it remains unclear how far predictions can be expanded, especially in species without known MHC-I ligands.

In this study, we introduce a pan-allele version of MixMHCpred (v3.0) trained on a large collection of MHC-I ligands across multiple species. This model directly bases its predictions on binding specificities which enabled us to carefully benchmark the accuracy of MHC-I ligand predictions across alleles and species. Our results demonstrate high accuracy of MHC-I ligand predictions across the vast majority of MHC-I alleles in human and in laboratory mouse strains, but reveal important limitations in species without experimentally determined MHC-I ligands.

## Results

### MHC-I peptidomics data reveal binding motifs and peptide length distributions for 143 MHC-I alleles

To expand the allelic coverage of our previous collection of MHC-I peptidomics datasets covering 119 human MHC-I alleles (Gfeller, Schmidt, et al., 2023), we collected data from several recent MHC-I peptidomics studies (DeVette et al., 2018; Ebrahimi-Nik et al., 2019; Faridi et al., 2020; Lampen et al., 2013; Marcu et al., 2021; Murphy et al., 2020; Nielsen et al., 2018; Pyke et al., 2021; Vita et al., 2019). These studies encompass MHC-I ligands from human, mouse, cattle, canid and non-human primate alleles. Motif deconvolution was performed on all samples following our previously established procedure (Bassani-Sternberg et al., 2017; Gfeller et al., 2018a; Gfeller, Schmidt, et al., 2023) (see Materials and Methods). This approach ultimately yielded a collection of 511,553 ligands (Figure 1A) interacting with 143 MHC-I alleles (Figure 1B, Suppl. Figure S1). As expected, the vast majority of ligands and alleles came from human (Figure 1A and B). For each allele, binding motifs (mathematically represented with position weight matrices) and peptide length distributions were computed (see Materials and Methods). Distinct motifs were built for each peptide length, ranging from 8- to 14-mers (Figure 1C), as ligands of varying lengths exhibit differences in their motifs.

**Figure 1:**
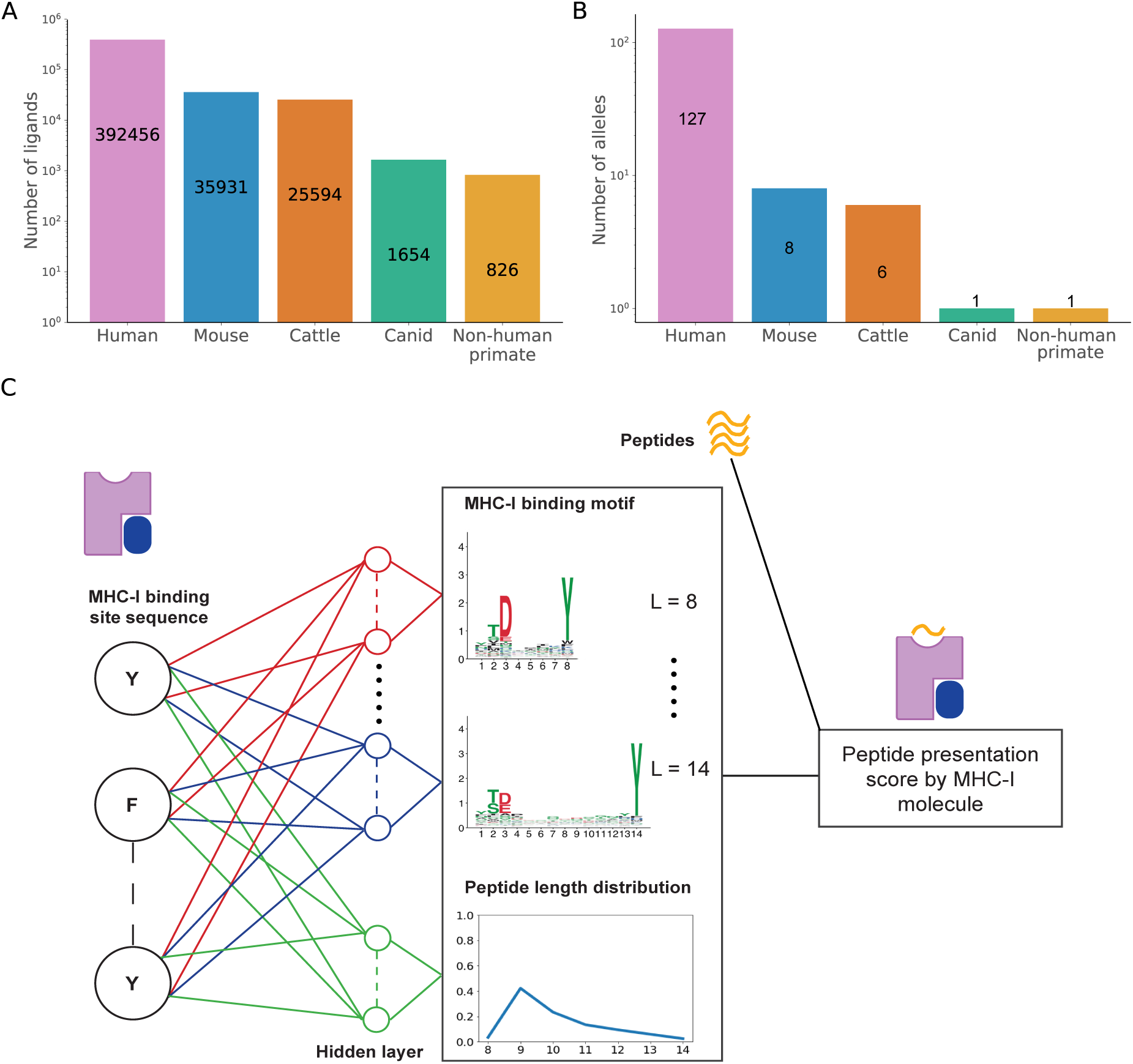
Overview of the MHC-I peptidomics data and model architecture used in our pan-allele MHC-I ligand predictor (MixMHCpred3.0). (A) Number of MHC-I ligands for MHC-I molecules from different species collected in this work. (B) Number of MHC-I alleles with known ligands from different species. (C) Description of the proposed architecture for predicting binding motifs and peptide length distribution (middle) and predicting peptide presentation scores (right). For the binding motif predictor, distinct neural networks were trained for each peptide length (from 8 to 14). An additional neural network was trained to predict peptide length distributions for MHC-I molecules without experimental ligands. The outputs of both predictors are combined in a final step to predict peptide presentation score and derive a %rank, as in previous versions of MixMHCpred.

### Binding motifs and peptide length distributions can be used to train a pan-allele MHC-I ligand predictor

We used the binding motifs and peptide length distributions to train a pan-allele predictor of MHC-I ligands, referred to as MixMHCpred3.0. Building upon our recent work (Racle et al., 2023; Tadros et al., 2023), we first trained neural networks to predict binding motifs for each peptide length, as well as peptide length distributions (Figure 1C). These networks use as input the MHC-I binding site sequence (see Materials and Methods). In a second step, MixMHCpred3.0 integrates the output from these neural networks and computes a final presentation score and %rank of a peptide (Figure 1C). This last step is performed following the procedure introduced for MixMHCpred2.2 (Gfeller, Schmidt, et al., 2023). This framework enables us to predict ligands for any MHC-I allele. In addition, it can accommodate either predicted motifs or motifs directly computed from experimental ligands for alleles with such data.

To test how reliably MHC-I ligand predictions can be extrapolated to MHC-I alleles without known ligands, we performed an extensive leave-one-allele-out (LOA) cross-validation. All ligands for each allele were iteratively excluded from the training of our model and used as test set, and the model was trained on the motifs and length distributions from the remaining alleles (see Materials and Methods). 99-fold excess of random peptides from the human proteome were used as negatives to compute the Area Under the Curve (AUC) of the Receiver Operating Characteristic Curve (ROC) values. These AUC values serve as an indicator of the model’s predictive power, with a value 1 for a perfect predictor and 0.5 in the case of random predictions. Overall, the predictions were much better than random for all alleles (Figure 2A). Predictions for human alleles also outperformed predictions for alleles in other species.

**Figure 2:**
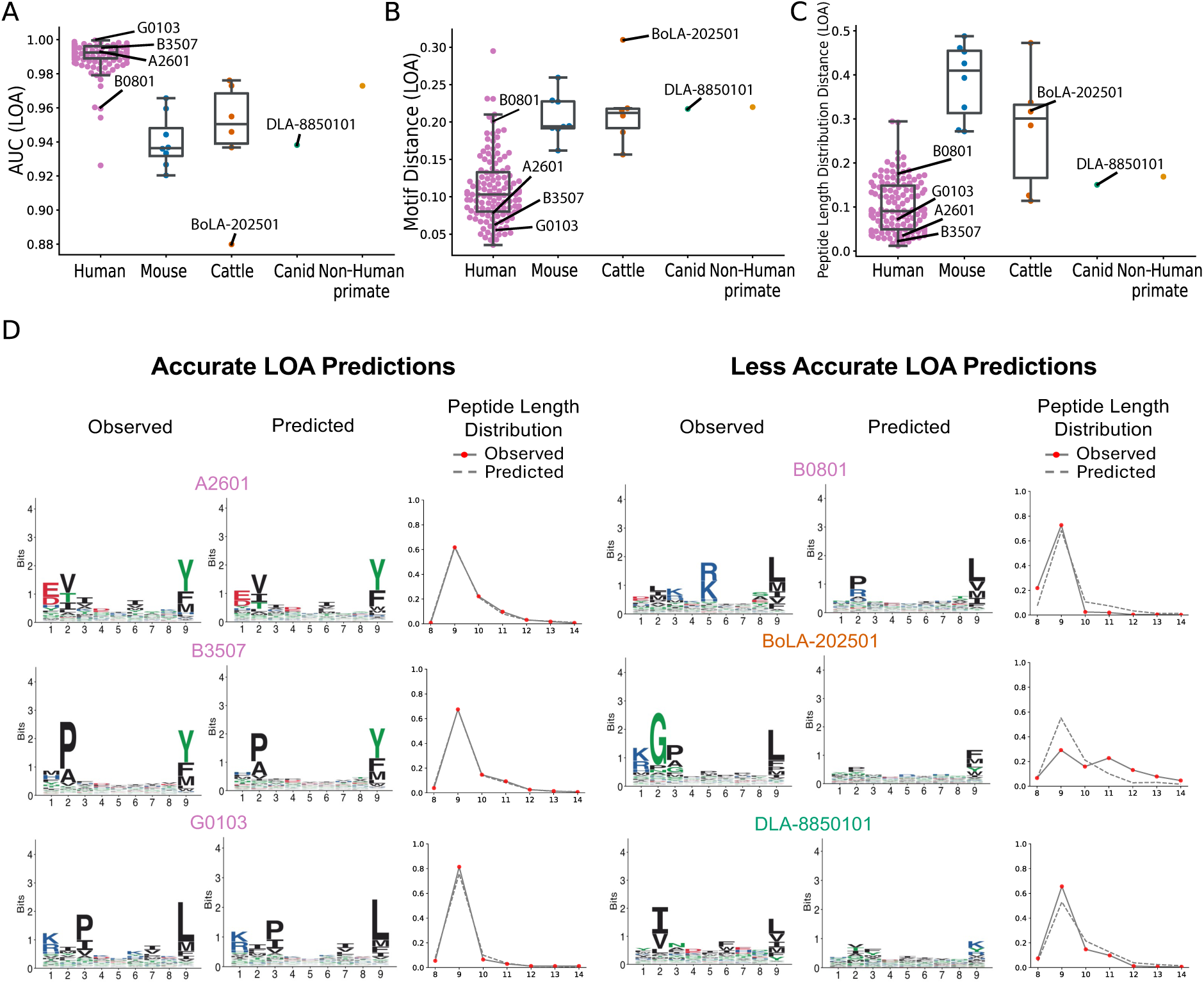
MHC-I binding specificity predictions for alleles without known ligands. (A) AUC for predictions of peptides presented by MHC-I molecules from different species, obtained in the leave-one-allele-out (LOA) cross-validation. (B) Euclidean distance between the predicted and experimental 9-mer motifs in the LOA cross-validation. (C) Euclidean distance between the predicted and experimental peptide length distributions in the LOA cross-validation. (D) Examples of predicted MHC-I binding motifs and peptide length distributions in a LOA context. Examples of 3 alleles with accurate predictions are shown on the left, and 3 alleles with less accurate predictions are shown on the right.

We then checked the accuracy of the predicted motifs and predicted length distributions, as an alternative to AUC for benchmarking. To this end, we computed the Euclidean distance between the predicted and actual 9-mer motifs (Figure 2B, see Materials and Methods). For most human alleles, the predicted binding motifs were highly similar to the actual ones. In other species, the distances between predicted and actual motifs were higher. Similar observations were made when comparing the Euclidean distance between predicted and experimental peptide length distributions (Figure 2C). These results demonstrate that both binding motifs and peptide length distributions for MHC-I alleles without known ligands can be accurately predicted in human, and less so in other species, thereby providing a rational explanation for the LOA AUC values in Figure 2A. To further visualize the quality of the model predictions and better interpret binding motif and peptide length distribution distances, we selected three alleles with very high LOA AUC values and three alleles with much lower LOA AUC values (Figure 2A). Figure 2D shows the comparison between the actual and the predicted 9-mer binding motifs and peptide length distributions. For alleles displaying very high LOA AUC (>0.98), we observed that binding motifs and peptide length distributions were indeed accurately predicted. For alleles with lower LOA AUC (0.88 to 0.95), we observed that the predicted binding motifs were less well predicted (Euclidean Distances of 0.2 to 0.3) and much less specific, capturing mainly a weakly conserved specificity for hydrophobic residues at the last position (Figure 2D). Similar observations could be made for peptide length distributions where the predicted peptide length distributions for alleles with lower LOA AUC were less accurate, while still capturing the preference for 9-mers (Figure 2D). These results also indicate that fairly high LOA AUC values of up to 0.95 can be obtained with relatively unspecific motifs. This likely reflects the fact that many MHC-I alleles share some similarity (e.g., preference for 9-mers, preference for hydrophobic residues at the last position of their ligands), which leads to some predictive power even when failing to capture the actual specificity of each allele. Accurately predicted motifs and peptide length distributions corresponded to cases with LOA AUC values around 0.98 or higher.

### Binding site similarity determines prediction accuracy

To explore the determinants of MHC-I ligand prediction accuracy for an allele without known ligands, we investigated the binding site similarity with alleles with known ligands. This binding site similarity was computed as a binding site sequence distance (see Materials and Methods). For each allele, we identified the closest allele with known ligands in terms of binding site distance and refer to this distance to the closest allele with known ligands as the “binding site distance”, for simplicity. We observed a strong inverse-correlation between binding site distances and AUC values computed in the LOA cross-validation (Figure 3A). As a rule of thumb, our data suggest that accurate MHC-I ligand predictions can be achieved when an allele shows a binding site distance smaller than 0.1 with another allele with known ligands. For bigger distances, predictions will generally be of lower accuracy.

**Figure 3:**
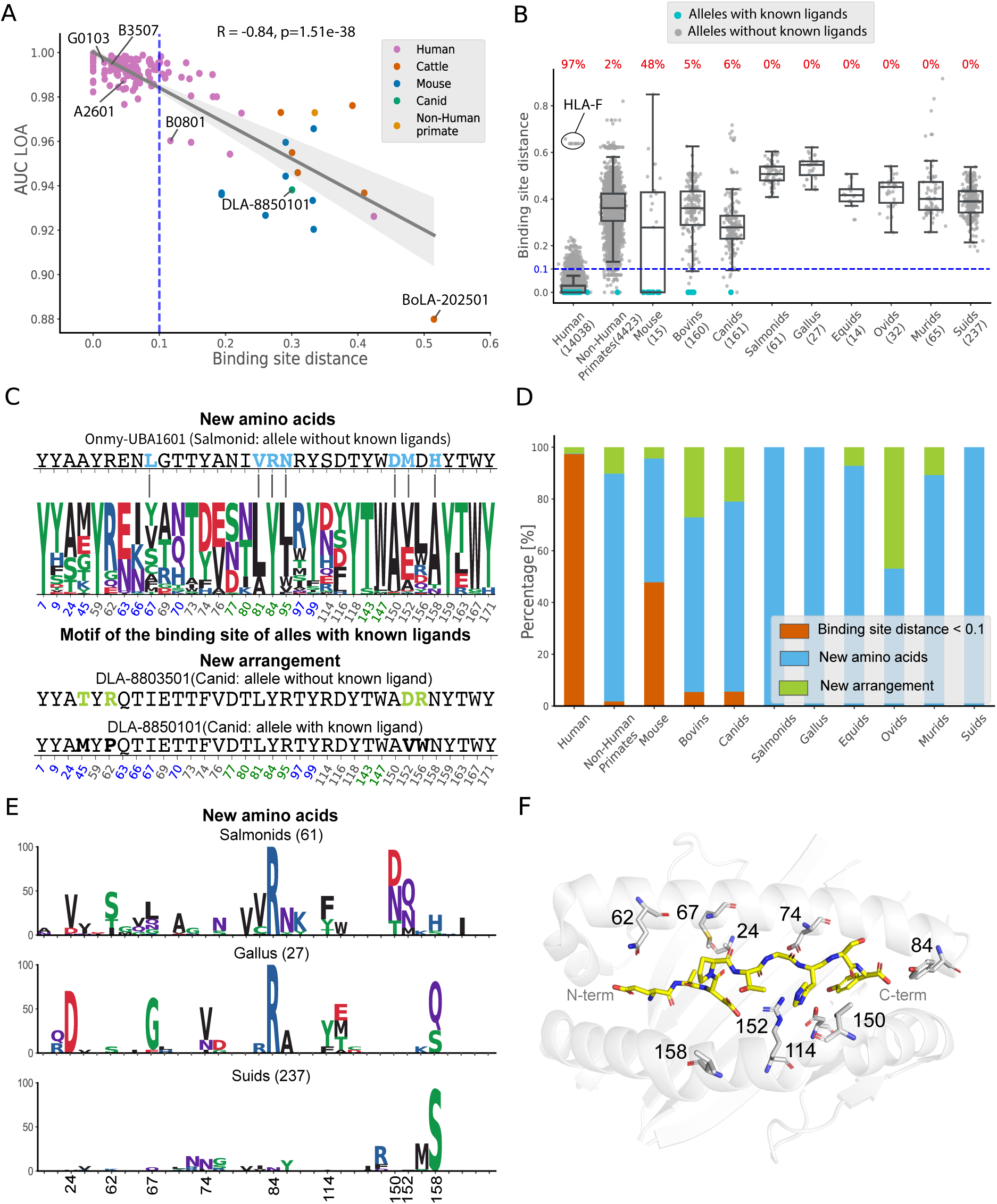
Binding site similarity determines prediction accuracy. (A) Relationship between the accuracy of MHC-I ligand predictions (AUC in the LOA cross-validation) and binding site distance to the closest allele with known ligands. Regression line and Pearson correlation coefficients were added to the plot. (B) Boxplots of binding site distances to the closest allele with known ligands for all known alleles in different species groups. The numbers above each boxplot show the percentage of MHC-I sequences in each group reaching a binding site distance lower than 0.1 (the blue dashed line). Numbers in parentheses indicate the total number of MHC-I alleles with available sequence in each species group. (C) Examples of different scenario characterizing alleles with binding site distances larger than 0.1. The amino acid frequency for the binding site positions for alleles with known ligands is shown in the middle. An example of an allele without known ligands and having new amino acids (i.e., unseen among alleles with known ligands, written in light blue) in its binding site is shown above. An example of an allele without ligands and having a different arrangement of amino acids is shown below (with amino acids non-conserved in its closest allele with known ligands indicated in yellow). B-pocket positions are marked in dark blue and F-pocket positions in green. (D) Stacked barplots showing the percentage of alleles with binding site distance < 0.1 (orange), alleles with binding site distance >= 0.1 and new amino acids at some binding site positions (light blue) and alleles with binding site distance >= 0.1 and new arrangements of amino acids in their binding site (yellow). (E) Frequency of the new amino acids in species where all MHC-I alleles have new amino acids compared to MHC-I alleles with known ligands (i.e., Salmonids, Gallus and Suids). (F) Representative 3D structure of the MHC-I binding site (HLA-A*01:01 in grey in complex with EADPTGHSY in yellow, PDB: 1W72), highlighting several positions with low conservation across species (see panel E).

We then investigated to which extent predictions with MixMHCpred3.0 could be applied with high confidence across all known MHC-I alleles. To this end, we collected over 19,000 MHC-I protein sequences from human and other species, extracted the binding site sequence for each allele, and computed its binding site distance with the closest allele with known ligands (see Materials and Methods). We then calculated for various species the fraction of alleles with a binding site distance lower than 0.1 (Figure 3B). We observed that more than 97% of human HLA-I alleles passed this threshold and are therefore expected to be accurately predicted. The few alleles with binding site distances > 0.5 corresponded to HLA-F alleles, for which we did not have reliable ligands in our data. This suggests that the 127 human HLA-I alleles with available ligands provide a very good coverage of the specificity space of human MHC-I alleles, including most HLA-A, HLA-B and HLA-C alleles. For mouse alleles, we observed 48% of the MHC-I alleles met the threshold and these include all alleles from laboratory mouse strains, for which ligands are available. For other species, our data show that most alleles without MHC-I ligands do not pass the threshold on the binding site distance, suggesting that predictions of MHC-I ligands and motifs will be of lower accuracy.

To enhance our understanding of these limitations in predicting MHC-I ligands in non-human species, we explored two different scenarios underlying binding site distances larger than 0.1 (Figure 3C). In the first scenario (“New Amino Acids”), some binding site positions display amino acids that are never found among alleles with known ligands (Suppl. Figure 2). In these cases, predictions are complicated, since the training set of MixMHCpred3.0 does not contain information about these ‘unseen’ amino acids. In the second scenario (“New Arrangements”), we considered cases where all amino acids in the binding site are found in alleles with known ligands, but not in a single allele. Figure 3C shows examples of these two scenarios. In the first case (Onmy-UBA1601, from Salmonids), we observed that several amino acids in the binding site (i.e., L67, V81, R84, N95, D150, M152 and H158) were not found among the alleles with known ligands. In the second case, (DLA-8803501 from canids), we observed that, even if all amino acids in the binding site are observed in alleles with known ligands, the closest allele with known ligands had different amino acids at 4 binding site positions (i.e., 45, 62, 152, 156). In general, we observed that most alleles with binding site distances larger than 0.1 displayed new amino acids in their binding site across all non-human species (blue bars in Figure 3D), which likely explains why their binding motifs are difficult to predict. Figure 3E shows the frequency of the new amino acids in the MHC-I binding site in three species—Salmonids, Gallus, and Suids—where all alleles showed some amino acids absent in alleles with known ligands. This analysis shows for instance that all MHC-I alleles in Salmonids and Gallus have R84, while all alleles with known ligands in our training set had Y84. Figure 3F shows the structural location in the MHC-I binding site of these less conserved positions and confirms that new amino acids at these positions could alter MHC-I binding specificity. Overall, these analyses suggest a molecular basis for the lower prediction accuracy observed in non-human species.

### MixMHCpred3.0 improves MHC-I ligand predictions

We then benchmarked our predictions for alleles without known ligands with two widely used pan-allele methods, NetMHCpan4.1 (Reynisson et al., 2020) and MHCflurry2.0 (O’Donnell et al., 2020), and the recently introduced predictor BigMHC (Albert et al., 2023). To this end, we first retrieved all alleles absent from the training sets of NetMHCpan (30 alleles in total), MHCflurry (10 alleles in total), or BigMHC (31 alleles in total) (Suppl. Table S1). We then retrained MixMHCpred3.0 excluding iteratively each of these alleles and tested the accuracy of the predictions on the ligands of the left-out alleles. Our results indicate similar performance with NetMHCpan (Figure 4A), and improved performance compared to MHCflurry (Figure 4B) and BigMHC (Figure 4C).

**Figure 4:**
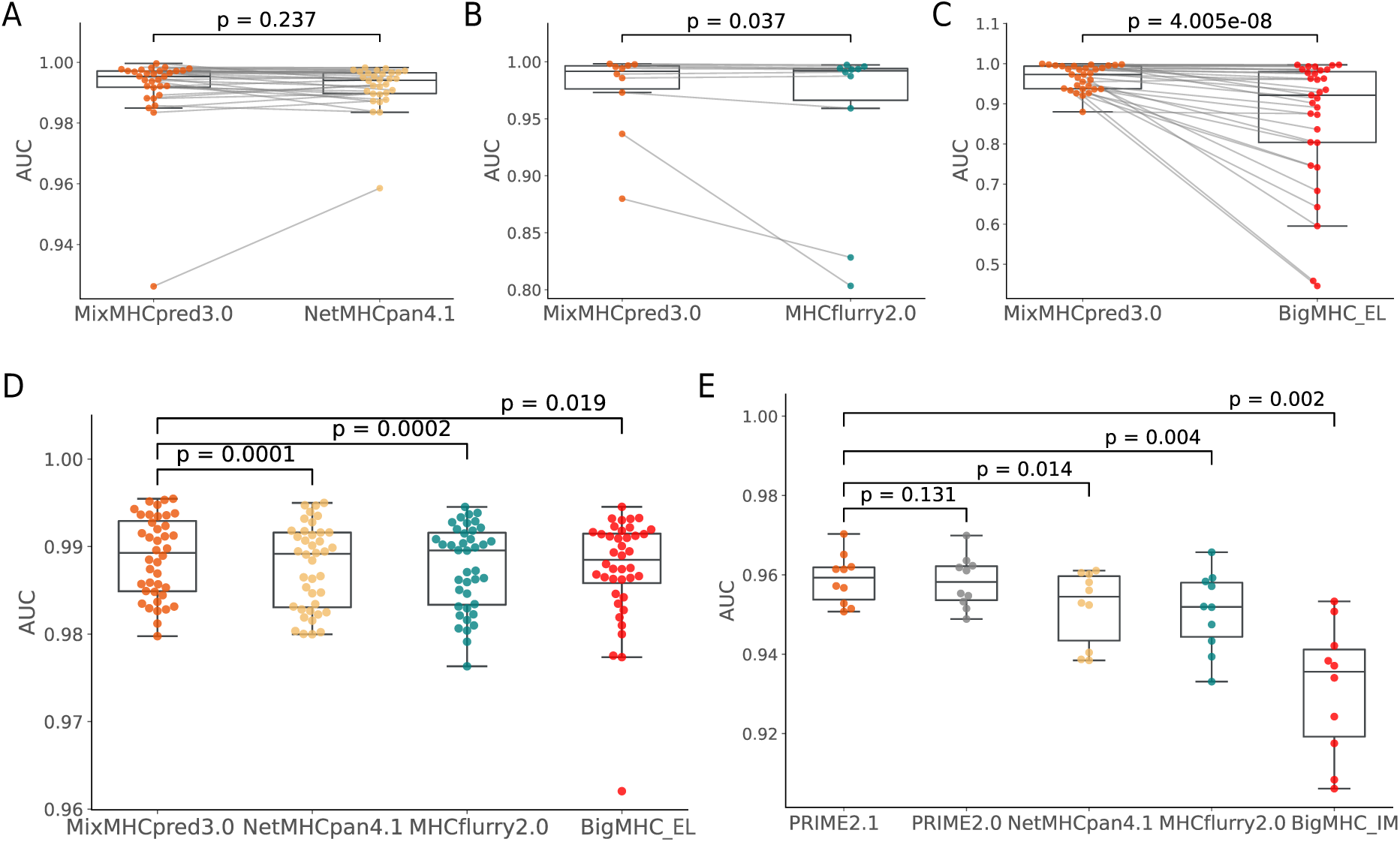
MixMHCpred3.0 compares favorably with other pan-allele MHC-I ligand predictors. (A-C) LOA cross-validation AUC values for the predictions of peptides presented by MHC-I molecules for alleles which are not part of the training set of (A) NetMHCpan, (B) MHCflurry or (C) BigMHC. (D) AUC values for the predictions of peptides presented by MHC-I coming from 40 samples in 3 different studies not included in the training set of any of the of the four predictors. (E) Benchmarking of PRIME2.1 on CD8+ T-cell epitope samples based on 10-fold cross-validation. P-values from a paired two-sided Wilcoxon signed rank test are indicated.

To further benchmark MixMHCpred3.0 on more realistic data, where most peptides come from alleles with known ligands, we employed three external datasets. The first dataset consists of ten HLA-I peptidomics samples from meningioma (Gfeller et al., 2018a), the second one includes ten HLA-I peptidomics samples (Pyke et al., 2021), and the third one comprises twenty recently published HLA-I peptidomics samples from COVID-19 (Nelde et al., 2022). To our knowledge, none of these datasets were used in the training of any predictor considered in this study. We employed 4-fold excess of random peptides from the human proteome as negatives to compute ROC curves (see Materials and Methods). The AUC values for MixMHCpred3.0 were significantly higher than those of NetMHCpan4.1, MHCflurry2.0 and BigMHC (Figure 4D). This demonstrates that MixMHCpred3.0 represents a state-of-the-art pan-allele MHC-I ligand predictor.

### MixMHCpred3.0 accurately predicts CD8+ T-cell epitopes

To ensure compatibility between MixMHCpred3.0 and our immunogenicity predictor PRIME (Gfeller, Schmidt, et al., 2023), we retrained PRIME with the scores provided by MixMHCpred3.0, resulting in the retrained version (PRIME2.1). To assess the impact of this integration on PRIME performance, we conducted a comprehensive benchmarking analysis. This analysis was designed to mirror the original validation methods described preciously in (Gfeller, Schmidt, et al., 2023) where we use CD8+ T-cell epitopes for benchmarking. The results of this analysis confirm the successful integration of MixMHCpred3.0 with PRIME, showing that the updated PRIME2.1 not only maintains compatibility with MixMHCpred3.0 but also surpasses the performance of other predictors (Figure 4E).

## DISCUSSION

CD8^+^ T-cell recognition of peptides displayed on MHC-I molecules plays a central role in immune recognition of infected or malignant cells. In this work, we capitalized on naturally presented MHC-I ligands derived from a diverse range of species, including human, mouse, cattle, canids, and non-human primate to train a pan-allele predictor of MHC-I ligands (MixMHCpred3.0) and explore how these predictions could be extrapolated across alleles and species.

Our results show that predictions of MHC-I ligands can be accurately expanded to MHC-I alleles with low binding site distance with respect to alleles with known ligands. These cases encompass the vast majority of human MHC-I alleles, indicating that pan-allele predictors are likely to work well even across individuals from diverse genetic backgrounds. The main exception consists of HLA-F alleles for which a consensus on their motifs has not been reached [PMID: **28636952 and 31717259**]. In other species, and especially in species with few known MHC-I ligands, predictions showed lower accuracy, and we expect also low accuracy for most other species without documented MHC-I ligands. This results from lower accuracy in predictions of both MHC-I binding motifs and peptides length distributions. We can attribute this limitation to the lower MHC-I binding site conservation in these species. In particular, the binding sites of MHC-I alleles from several species included amino acids which were never seen in alleles with known ligands.. These observations have implications for designing experiments aimed at improving the coverage of MHC-I alleles for which predictions of ligands can be made. In particular, we anticipate that MHC-I peptidomics profiling of alleles with binding site sequences including D24, S62, G67, R84, D/N150 or S/Q158 could reveal novel MHC-I binding motifs and expand our ability to accurately predict ligands for alleles across a broader range of species.

When performing our LOA cross-validation, we observed that AUC values up to 0.95 could be obtained even with relatively unspecific motifs (see examples in Figure 2D). This suggests that using only AUC as a performance metric does not guarantee that a pan-allele MHC-I ligand predictor has accurately learned the specificities of each allele. It also indicates that binding motifs and peptide length distributions, which are at the core of the pan-allele architecture of MixMHCpred3.0, provide a useful quality-control to evaluate the ability of a pan-allele MHC-I ligand predictor to learn the actual specificity of different alleles. For most practical applications in epitope discovery, the number of peptides which are being scored is much higher than the set of actual epitopes. Therefore very high AUC (e.g., >0.98) are desirable to have enough true positives among the top predicted peptides which are typically considered for experimental validation.

When comparing predictions of MixMHCpred3.0 for MHC-I alleles without known ligands (i.e., LOA cross-validation) with those generated by other methods such as NetMHCpan4.1, MHCflurry2.0 and BigMHC, we observed similar or better performance for MixMHCpred3.0. This suggests that our observations about the strengths and limitations to extrapolate predictions to alleles without known ligands will also apply to these other tools.

In most practical applications for epitope discovery, MHC-I ligand predictions are performed on human samples where the majority of MHC-I alleles have known ligands in existing databases. The improved predictions of MixMHCpred3.0 compared with other tools in such cases indicate that MixMHCpred3.0 provides a state-of-the-art solution with high computational efficiency. Moreover, the compatibility with the PRIME framework (Gfeller, Schmidt, et al., 2023) ensures that MixMHCpred3.0 can be used for CD8^+^ T-cell epitope discovery.

Altogether, our work provides an integrated pan-allele version of the MixMHCpred tool (https://github.com/GfellerLab/MixMHCpred). This new version expands the allelic coverage of MixMHCpred and preserves its distinctive interpretability in terms of binding motifs and peptide length distribution for different alleles. Our careful validation demonstrates that accurate and specific predictions can be achieved for the vast majority of human MHC-I alleles and for mouse MHC-I alleles from laboratory mouse strains. Our results further highlight the importance of expanding experimental characterization of binding motifs for alleles in other species to reach reliable MHC-I ligand predictions across a broader range of species.

## MATERIALS AND METHODS

### Collection of MHC-I ligands

Naturally presented MHC-I ligands were collected from more than 250 MHC-I peptidomics samples from human, mouse, cattle, canid, and non-human primate. These include all samples considered in (Gfeller, Schmidt, et al., 2023). We further included data from a few recent MHC-I peptidomics studies (DeVette et al., 2018; Ebrahimi-Nik et al., 2019; Faridi et al., 2020; Lampen et al., 2013; Marcu et al., 2021; Murphy et al., 2020; Nielsen et al., 2018; Pyke et al., 2021; Vita et al., 2019). All data were retrieved from the original studies to prevent having filtered data based on MHC-I ligand predictors. All samples were processed with our motif deconvolution tool (MixMHCp) to identify shared motifs across samples sharing the same allele (Gfeller et al., 2018a). Further information regarding this procedure and the results obtained can be found in our previous publications (Bassani-Sternberg et al., 2017b; Gfeller et al., 2018a; Gfeller, Schmidt, et al., 2023). The final dataset of naturally presented MHC-I ligands comprises 511,553 peptide-MHC-I interactions with 143 different MHC-I alleles.

### Building MHC-I binding motifs and peptide length distributions

For all MHC-I alleles with naturally presented ligands, Position Probability Matrices (PPMs) were constructed by computing the frequency of each amino acid at each position in the set of ligands of the given allele, including standard pseudocounts based on BLOSUM62 as detailed in (Gfeller et al., 2018 and Racle et. al, 2019). Separate PPMs were generated for each ligand length L from 8 to 14. The Position Weight Matrices representing the final binding motifs were computed by normalizing the PPMs with the amino acid background frequencies of the human proteome, as outlined in (Gfeller et al., 2018a; Racle et al., 2019). Binding motifs were visualized using ggseqlogo (Wagih, 2017) and Logomaker (Tareen & Kinney, 2020). To determine peptide length distributions, the fraction of naturally presented MHC-I ligands of each length (from 8 to 14) was computed as described in (Gfeller et al., 2018a).

### Predicting MHC-I binding motifs

Inspired by our recent work on MHC-I and MHC-II motifs (Racle et al., 2023; Tadros et al., 2023), neural networks were used to predict PPMs of MHC-I molecules without known ligands. More precisely, distinct networks were trained for each peptide length (8 to 14). The input of each neural network is the list of binding site residues from the MHC-I molecules (34 residues). This binding site was defined as in (Hoof et al., 2009). Each binding site residue was encoded as a 20-dimensional vector based on the BLOSUM62 probability matrix. The output of each network consists of a matrix of 20xL, representing the PPM at the corresponding motif length L. Each network is composed of an input layer (34×20 nodes), one fully connected hidden layer (256 nodes) followed by a dropout of 0.2 and an additional layer that reshapes the output layer from a vector (20xL nodes) to a matrix of size 20 rows and L columns. We used rectified linear unit (ReLU) activation function for the hidden layer and a custom softmax function for the output layer that applies the softmax activation function on each column of the matrix. We used the Kullback Leibler divergence as a loss function, and it was optimized using Adam optimizer with a learning rate of 0.0001. These neural networks were implemented in Python (version 3.7.11), using Keras packages relying on TensorFlow (version 2.2.4-tf). 1000 epochs were set for the training process. For each allele and each peptide length (8 to 14), we then normalized by background human proteome frequencies to create the final predicted PWM.

### Predicting peptide length distributions

A neural network was developed to predict the peptide length distribution of MHC-I molecules. The input layer is the same as for the MHC-I motifs prediction (34×20 nodes), followed by one hidden layer (128 nodes) with the rectified linear unit (ReLU) activation function followed by a dropout of 0.2. The output layer is the peptide length distribution (from 8 to 14, i.e., 7 nodes) based on the softmax activation function. We used the Kullback Leibler divergence as a loss function, and it was optimized using Adam optimizer with a learning rate of 0.0001. A maximum of 125 epochs were set for the training process with early stopping applied if no improvement in loss was observed over a span of 20 consecutive epochs.

### Leave-one-allele-out (LOA) cross-validation

We performed leave-one-allele-out cross-validation for ligand, binding motif and peptide length distribution predictions using iteratively as test set each allele. For ligand predictions, a 99-fold excess of negatives were added randomly from the human proteome with uniform length distribution from 8 to 14. Subsequently, binding scores were predicted for each peptide in the test set, and the performance was evaluated based on the AUC values (Figure 2A). For each length (from 8 to 14), the predicted PWMs were compared to the experimental ones by computing the Euclidean distance for each position of the PWM and averaging these distances. The lower the distance, the closer the predicted motifs are to the experimental ones (Figure 2B shows the Euclidean distance for the 9-mers motifs). Similarly, we computed the Euclidean distance between the predicted and experimental peptide length distributions (Figure 2C).

#### Binding site sequence distances

The binding site distance between two alleles was calculated as described in the following formula:

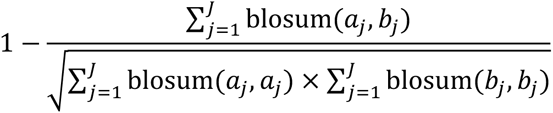

in which blosum refers to the Blosum62 scoring matrix, which is used to score amino acid substitutions (Henikoff & Henikoff, 1992), J represents the length of the MHC-I binding site sequence (34 amino acids), a_j_ and b_j_ denote the amino acid from the two alleles being compared. The resulting score ranges from 0 to 1, where a higher score indicates a greater distance between the two MHC-I binding site sequences. For alleles without known ligands, the binding site distance to the set of alleles with known ligands is defined as the minimum of the distances to alleles with known ligand, or equivalently the distance to the closest alleles with known ligands. For simplicity, this distance to the closest allele is often referred to as the “binding site distance”.

#### MHC-I sequences retrieval and alignment

Human MHC-I sequences were retrieved from IPD-IMGT/HLA database (Barker et al., 2023). MHC-I sequences from multiple other species were retrieved from IPD-MHC database (Maccari et al., 2017). Mouse MHC-I sequences are not part of IPD-MHC database so they were manually retrieved from the UniProtKB database (The UniProt Consortium et al., 2021). We then aligned the MHC-I sequences using the MAFFT algorithm (Katoh, 2002, version 7.520) and took as reference the list of sequences of alleles with known ligands for the alignment.

#### Benchmarking with other MHC-I ligand predictors

To assess the accuracy of the ligand predictions for MHC-I molecules and compare with the state-of-the-art methods, such as NetMHCpan, MHCflurry and BigMHC, we performed the leave-one-allele-out cross-validation, where each allele absent from the training of NetMHCpan4.1, MHCflurry2.0 and BigMHC was successively removed from the training set of MixMHCpred3.0 (30, 10 and 31 alleles respectively) (see Figures 4A, B and first, Suppl. Table S1).

In the second benchmark based on full HLA-I peptidomics samples, we used HLA-I peptidomics datasets coming from 10 menigioma samples measured in (Gfeller et al., 2018a) and 10 HLA-I peptidomics samples from (Pyke et al., 2021) that were not part of the training of any version of MixMHCpred, NetMHCpan, MHCflurry nor BigMHC. To these, we added a third dataset that comprises twenty recently published HLA-I peptidomics samples from COVID-19 patients (Nelde et al., 2022). In their paper, the full HLA-I typing was not provided for each sample, so we ran our motif deconvolution tool (MixMHCp) (Gfeller et al., 2018a) to annotate the alleles to 4 digits typing in each sample excluding sample ‘UPN17’ due to ambiguity in HLA-I annotation. All peptides from a given sample were used together with the set of alleles describing this sample and considered as positives and we added four times more random peptides as negatives. The scores for all peptides across all alleles were calculated, and the best score among the alleles of a sample was retained (%Rank_bestAllele for MixMHCpred, lowest %Rank_EL for NetMHCpan, presentation_percentile for MHCflurry, highest BigMHC_EL score for BigMHC). Using the predicted scores for each peptide, the AUC was computed separately for each predictor and sample, providing a performance evaluation and comparison with existing state-of-the-art methods (see Figure 4D).

#### PRIME2.1 benchmarking

PRIME was retrained with the scores provided by MixMHCpred3.0, resulting in the updated version, PRIME2.1. The benchmarking of PRIME2.1 based on 10-fold cross-validation presented in Figure 4E used exactly the same data as in PRIME2.0 publication (Gfeller, Schmidt, et al., 2023).

## ACKNOWLEDGMENTS

We thank Matei Teleman and Aurelie AG Gabriel for testing the MixMHCpred tool.

## CONTRIBUTIONS

D.T. performed the new methodological developments, wrote the manuscript and prepared the figures. J.R. provided feedback for the project and the manuscript. D.G. designed the project, supervised the work and provided feedback for the manuscript.

## DATA & CODE AVAILABILITY

- MixMHCpred3.0 and PRIME2.1 are available at https://github.com/GfellerLab/
- Any additional information required to reproduce this work is available from the Lead Contact upon request.

## Supplementary Materials

**Suppl. Table S1:** Alleles with known ligands absent from the training sets of NetMHCpan (30 alleles in total), MHCflurry (10 alleles in total), or BigMHC (31 alleles in total).

| NetMHCpan4.1 | MHCflurry2.0 | BigMHC |
| --- | --- | --- |
| A0252 A1102 A2407 | A0220 A0252 A2608 | A0220 A0252 A2608 |
| A2608 A3401 A3402 | B1805 B4032 B6701 | B1401 B1805 B4032 |
| A3601 A7401 B0704 | C1602 BoLA-102301 | B4801 B6701 B8101 |
| B1513 B1805 B3507 | BoLA-202501 | C1505 C1602 E0103 |
| B3802 B3905 B4006 | BoLA-300201 | G0101 G0103 G0104 |
| B4032 B4405 B5502 |  | H2-Db H2-Dd H2-Dq |
| B5802 B8101 C0302 |  | H2-Kb H2-Kd H2-Kk |
| C0403 C0704 C1204 |  | H2-Kq H2-Ld |
| C1403 C1602 E0103 |  | BoLA-102301 |
| G0101 G0103 G0104 |  | BoLA-201201 |
|  |  | BoLA-202501 |
|  |  | BoLA-300201 |
|  |  | BoLA-402401 |
|  |  | BoLA-601301 |
|  |  | DLA-8850101 |
|  |  | Mamu-B00801 |

**Suppl. Figure S1:**
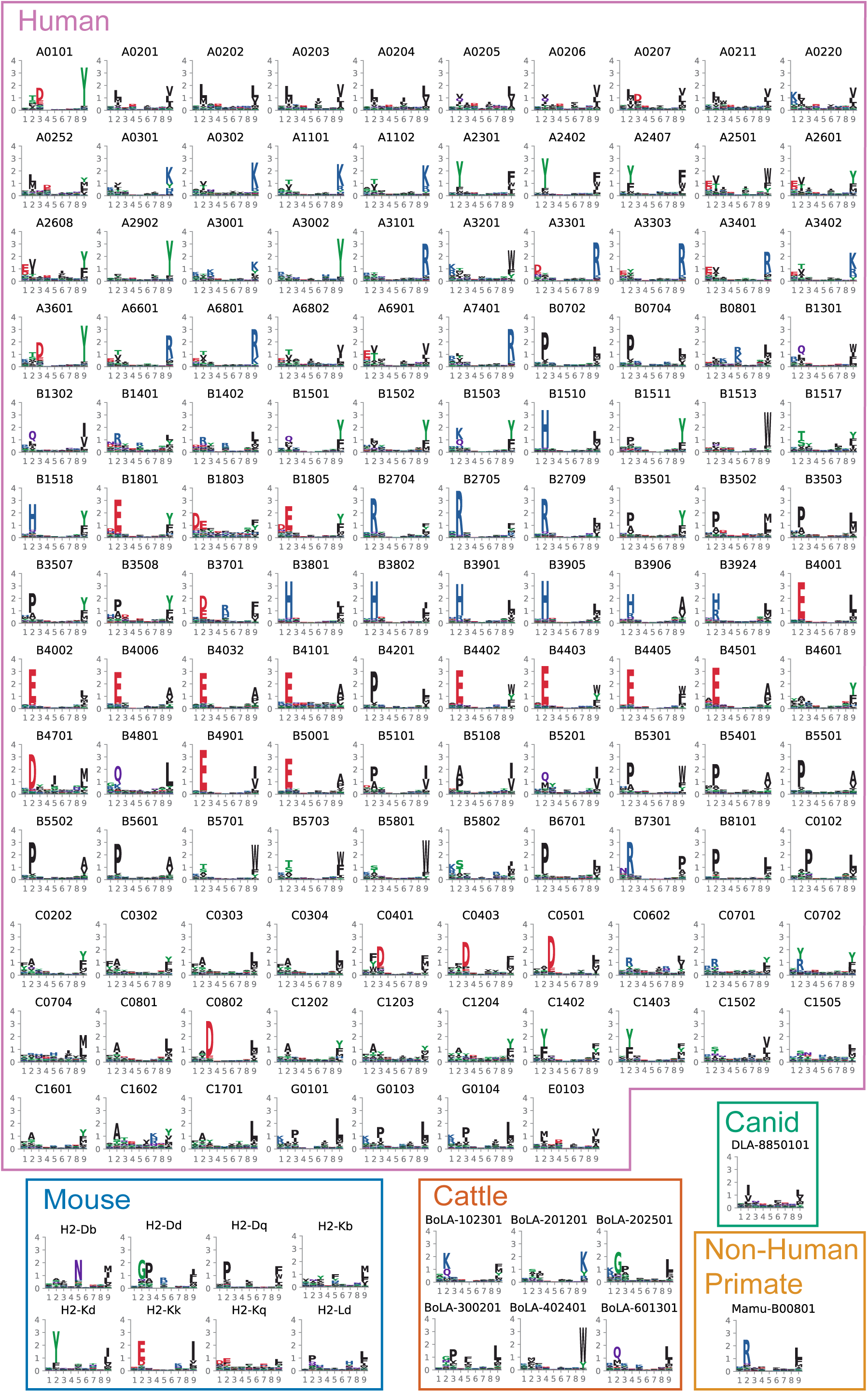
Curation of MHC-I peptidomics data reveals binding specificities for 143 MHC-I alleles, related to. Figure 1: 9-mers MHC binding motifs describing the binding specificities of the 143 MHC-I alleles determined in this study.

**Suppl. Figure S2:**
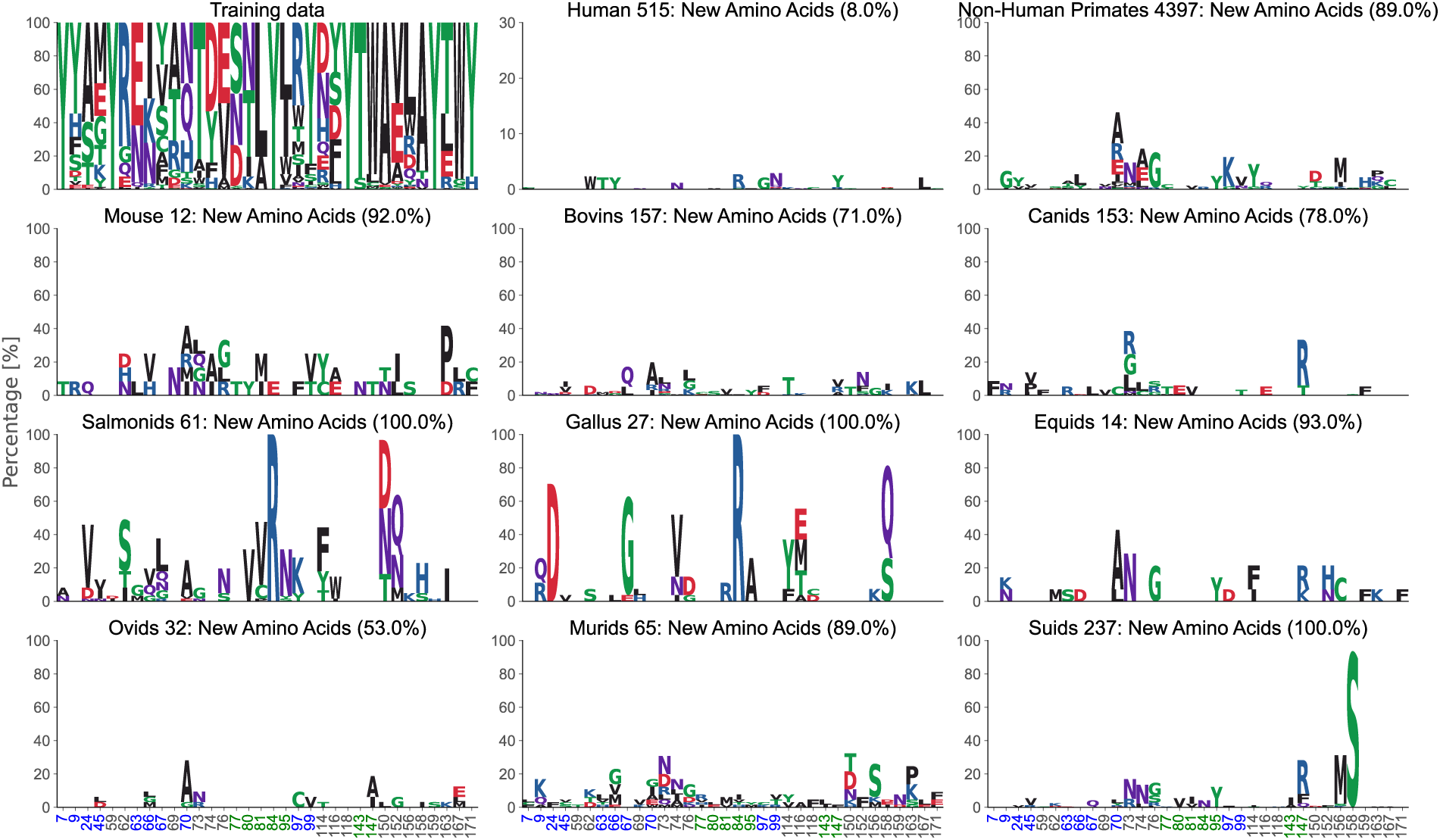
New amino acids for alleles without known ligands, related to. Figure 3: Binding site sequence motifs of alleles with known ligands (training data, upper left) and the frequency of the new amino acids in species where all MHC-I alleles have new amino acids compared to MHC-I alleles with known ligands.

## REFERENCES

Abelin, J. G., Keskin, D. B., Sarkizova, S., Hartigan, C. R., Zhang, W., Sidney, J., Stevens, J., Lane, W., Zhang, G. L., Eisenhaure, T. M., Clauser, K. R., Hacohen, N., Rooney, M. S., Carr, S. A., & Wu, C. J. (2017). Mass Spectrometry Profiling of HLA-Associated Peptidomes in Mono-allelic Cells Enables More Accurate Epitope Prediction. Immunity, 46(2), 315–326. 10.1016/j.immuni.2017.02.007

Albert, B. A., Yang, Y., Shao, X. M., Singh, D., Smith, K. N., Anagnostou, V., & Karchin, R. (2023). Deep neural networks predict class I major histocompatibility complex epitope presentation and transfer learn neoepitope immunogenicity. Nature Machine Intelligence, 5(8), 861–872. 10.1038/s42256-023-00694-6

Andreatta, M., & Nielsen, M. (2016). Gapped sequence alignment using artificial neural networks: Application to the MHC class I system. Bioinformatics, 32(4), 511–517. 10.1093/bioinformatics/btv639

Barker, D. J., Maccari, G., Georgiou, X., Cooper, M. A., Flicek, P., Robinson, J., & Marsh, S. G. E. (2023). The IPD-IMGT/HLA Database. Nucleic Acids Research, 51(D1), D1053– D1060. 10.1093/nar/gkac1011

Bassani-Sternberg, M., Chong, C., Guillaume, P., Solleder, M., Pak, H., Gannon, P. O., Kandalaft, L. E., Coukos, G., & Gfeller, D. (2017a). Deciphering HLA-I motifs across HLA peptidomes improves neo-antigen predictions and identifies allostery regulating HLA specificity. PLOS Computational Biology, 13(8), e1005725. 10.1371/journal.pcbi.1005725

Bassani-Sternberg, M., Chong, C., Guillaume, P., Solleder, M., Pak, H., Gannon, P. O., Kandalaft, L. E., Coukos, G., & Gfeller, D. (2017b). Deciphering HLA-I motifs across HLA peptidomes improves neo-antigen predictions and identifies allostery regulating HLA specificity. PLoS Computational Biology, 13(8), e1005725. 10.1371/journal.pcbi.1005725

Bernatchez, L., & Landry, C. (2003). MHC studies in nonmodel vertebrates: What have we learned about natural selection in 15 years? Journal of Evolutionary Biology, 16(3), 363–377. 10.1046/j.1420-9101.2003.00531.x

Bulik-Sullivan, B., Busby, J., Palmer, C. D., Davis, M. J., Murphy, T., Clark, A., Busby, M., Duke, F., Yang, A., Young, L., Ojo, N. C., Caldwell, K., Abhyankar, J., Boucher, T., Hart, M. G., Makarov, V., De Montpreville, V. T., Mercier, O., Chan, T. A., … Yelensky, R. (2019). Deep learning using tumor HLA peptide mass spectrometry datasets improves neoantigen identification. Nature Biotechnology, 37(1), 55–63. 10.1038/nbt.4313

Carreno, B. M., Magrini, V., Becker-Hapak, M., Kaabinejadian, S., Hundal, J., Petti, A. A., Ly, A., Lie, W.-R., Hildebrand, W. H., Mardis, E. R., & Linette, G. P. (2015). A dendritic cell vaccine increases the breadth and diversity of melanoma neoantigen-specific T cells. Science, 348(6236), 803–808. 10.1126/science.aaa3828

DeVette, C. I., Andreatta, M., Bardet, W., Cate, S. J., Jurtz, V. I., Jackson, K. W., Welm, A. L., Nielsen, M., & Hildebrand, W. H. (2018). NetH2pan: A Computational Tool to Guide MHC Peptide Prediction on Murine Tumors. Cancer Immunology Research, 6(6), 636–644. 10.1158/2326-6066.CIR-17-0298

Ebrahimi-Nik, H., Michaux, J., Corwin, W. L., Keller, G. L. J., Shcheglova, T., Pak, H., Coukos, G., Baker, B. M., Mandoiu, I. I., Bassani-Sternberg, M., & Srivastava, P. K. (2019). Mass spectrometry–driven exploration reveals nuances of neoepitope-driven tumor rejection. JCI Insight, 4(14), e129152. 10.1172/jci.insight.129152

Faridi, P., Woods, K., Ostrouska, S., Deceneux, C., Aranha, R., Duscharla, D., Wong, S. Q., Chen, W., Ramarathinam, S. H., Lim Kam Sian, T. C. C., Croft, N. P., Li, C., Ayala, R., Cebon, J. S., Purcell, A. W., Schittenhelm, R. B., & Behren, A. (2020). Spliced Peptides and Cytokine-Driven Changes in the Immunopeptidome of Melanoma. Cancer Immunology Research, 8(10), 1322–1334. 10.1158/2326-6066.CIR-19-0894

Gfeller, D., & Bassani-Sternberg, M. (2018). Predicting Antigen Presentation—What Could We Learn From a Million Peptides? Frontiers in Immunology, 9, 1716. 10.3389/fimmu.2018.01716

Gfeller, D., Guillaume, P., Michaux, J., Pak, H.-S., Daniel, R. T., Racle, J., Coukos, G., & Bassani-Sternberg, M. (2018a). The Length Distribution and Multiple Specificity of Naturally Presented HLA-I Ligands. Journal of Immunology (Baltimore, Md.: 1950), 201(12), 3705–3716. 10.4049/jimmunol.1800914

Gfeller, D., Guillaume, P., Michaux, J., Pak, H.-S., Daniel, R. T., Racle, J., Coukos, G., & Bassani-Sternberg, M. (2018b). The Length Distribution and Multiple Specificity of Naturally Presented HLA-I Ligands. The Journal of Immunology, 201(12), 3705–3716. 10.4049/jimmunol.1800914

Gfeller, D., Liu, Y., & Racle, J. (2023). Contemplating immunopeptidomes to better predict them. Seminars in Immunology, 66, 101708. 10.1016/j.smim.2022.101708

Gfeller, D., Schmidt, J., Croce, G., Guillaume, P., Bobisse, S., Genolet, R., Queiroz, L., Cesbron, J., Racle, J., & Harari, A. (2023). Improved predictions of antigen presentation and TCR recognition with MixMHCpred2.2 and PRIME2.0 reveal potent SARS-CoV-2 CD8+ T-cell epitopes. Cell Systems, 14(1), 72–83.e5. 10.1016/j.cels.2022.12.002

Heitmann, J. S., Bilich, T., Tandler, C., Nelde, A., Maringer, Y., Marconato, M., Reusch, J., Jäger, S., Denk, M., Richter, M., Anton, L., Weber, L. M., Roerden, M., Bauer, J., Rieth, J., Wacker, M., Hörber, S., Peter, A., Meisner, C., … Walz, J. S. (2022). A COVID-19 peptide vaccine for the induction of SARS-CoV-2 T cell immunity. Nature, 601(7894), 617–622. 10.1038/s41586-021-04232-5

Henikoff, S., & Henikoff, J. G. (1992). Amino acid substitution matrices from protein blocks. Proceedings of the National Academy of Sciences, 89(22), 10915–10919. 10.1073/pnas.89.22.10915

Hoof, I., Peters, B., Sidney, J., Pedersen, L. E., Sette, A., Lund, O., Buus, S., & Nielsen, M. (2009). NetMHCpan, a method for MHC class I binding prediction beyond humans. Immunogenetics, 61(1), 1–13. 10.1007/s00251-008-0341-z

Hu, Y., Wang, Z., Hu, H., Wan, F., Chen, L., Xiong, Y., Wang, X., Zhao, D., Huang, W., & Zeng, J. (2019). ACME: Pan-specific peptide–MHC class I binding prediction through attention-based deep neural networks. Bioinformatics, 35(23), 4946–4954. 10.1093/bioinformatics/btz427

Katoh, K. (2002). MAFFT: A novel method for rapid multiple sequence alignment based on fast Fourier transform. Nucleic Acids Research, 30(14), 3059–3066. 10.1093/nar/gkf436

Lampen, M. H., Hassan, C., Sluijter, M., Geluk, A., Dijkman, K., Tjon, J. M., de Ru, A. H., van der Burg, S. H., van Veelen, P. A., & van Hall, T. (2013). Alternative peptide repertoire of HLA-E reveals a binding motif that is strikingly similar to HLA-A2. Molecular Immunology, 53(1–2), 126–131. 10.1016/j.molimm.2012.07.009

Leidner, R., Sanjuan Silva, N., Huang, H., Sprott, D., Zheng, C., Shih, Y.-P., Leung, A., Payne, R., Sutcliffe, K., Cramer, J., Rosenberg, S. A., Fox, B. A., Urba, W. J., & Tran, E. (2022). Neoantigen T-Cell Receptor Gene Therapy in Pancreatic Cancer. New England Journal of Medicine, 386(22), 2112–2119. 10.1056/NEJMoa2119662

Lundegaard, C., Lund, O., & Nielsen, M. (2008). Accurate approximation method for prediction of class I MHC affinities for peptides of length 8, 10 and 11 using prediction tools trained on 9mers. <u>Bioinformatics</u>, 24(11), 13971–398. 10.1093/bioinformatics/btn128

Maccari, G., Robinson, J., Ballingall, K., Guethlein, L. A., Grimholt, U., Kaufman, J., Ho, C.-S., de Groot, N. G., Flicek, P., Bontrop, R. E., Hammond, J. A., & Marsh, S. G. E. (2017). IPD-MHC 2.0: An improved inter-species database for the study of the major histocompatibility complex. Nucleic Acids Research, 45(D1), D860–D864. 10.1093/nar/gkw1050

Marcu, A., Bichmann, L., Kuchenbecker, L., Kowalewski, D. J., Freudenmann, L. K., Backert, L., Mühlenbruch, L., Szolek, A., Lübke, M., Wagner, P., Engler, T., Matovina, S., Wang, J., Hauri-Hohl, M., Martin, R., Kapolou, K., Walz, J. S., Velz, J., Moch, H., … Neidert, M. C. (2021). HLA Ligand Atlas: A benign reference of HLA-presented peptides to improve T-cell-based cancer immunotherapy. Journal for Immunotherapy of Cancer, 9(4), e002071. 10.1136/jitc-2020-002071

Migliorini, D., Dutoit, V., Allard, M., Grandjean Hallez, N., Marinari, E., Widmer, V., Philippin, G., Corlazzoli, F., Gustave, R., Kreutzfeldt, M., Blazek, N., Wasem, J., Hottinger, A., Koka, A., Momjian, S., Lobrinus, A., Merkler, D., Vargas, M.-I., Walker, P. R., … Dietrich, P.-Y. (2019). Phase I/II trial testing safety and immunogenicity of the multipeptide IMA950/poly-ICLC vaccine in newly diagnosed adult malignant astrocytoma patients. Neuro-Oncology, 21(7), 923– 933. 10.1093/neuonc/noz040

Murphy, J. P., Yu, Q., Konda, P., Paulo, J. A., Jedrychowski, M. P., Kowalewski, D. J., Schuster, H., Kim, Y., Clements, D., Jain, A., Stevanovic, S., Gygi, S. P., Mancias, J. D., & Gujar, S. (2020). Multiplexed Relative Quantitation with Isobaric Tagging Mass Spectrometry Reveals Class I Major Histocompatibility Complex Ligand Dynamics in Response to Doxorubicin. 19.

Neefjes, J., Jongsma, M. L. M., Paul, P., & Bakke, O. (2011). Towards a systems understanding of MHC class I and MHC class II antigen presentation. Nature Reviews Immunology, 11(12), 823–836. 10.1038/nri3084

Nelde, A., Rieth, J., Roerden, M., Dubbelaar, M. L., Hoenisch Gravel, N., Bauer, J., Klein, R., Hoheisel, T., Mahrhofer, H., Göpel, S., Bitzer, M., Hörber, S., Peter, A., Heitmann, J. S., & Walz, J. S. (2022). Increased soluble HLA in COVID-19 present a disease-related, diverse immunopeptidome associated with T cell immunity. iScience, 25(12), 105643. 10.1016/j.isci.2022.105643

Nielsen, M., & Andreatta, M. (2016). NetMHCpan-3.0; improved prediction of binding to MHC class I molecules integrating information from multiple receptor and peptide length datasets. Genome Medicine, 8(1), 33. 10.1186/s13073-016-0288-x

Nielsen, M., Connelley, T., & Ternette, N. (2018). Improved Prediction of Bovine Leucocyte Antigens (BoLA) Presented Ligands by Use of Mass-Spectrometry-Determined Ligand and in Vitro Binding Data. Journal of Proteome Research, 17(1), 559–567. 10.1021/acs.jproteome.7b00675

O’Donnell, T. J., Rubinsteyn, A., & Laserson, U. (2020). MHCflurry 2.0: Improved Pan-Allele Prediction of MHC Class I-Presented Peptides by Incorporating Antigen Processing. Cell Systems, 11(1), 42–48.e7. 10.1016/j.cels.2020.06.010

Ott, P. A., Hu, Z., Keskin, D. B., Shukla, S. A., Sun, J., Bozym, D. J., Zhang, W., Luoma, A., Giobbie-Hurder, A., Peter, L., Chen, C., Olive, O., Carter, T. A., Li, S., Lieb, D. J., Eisenhaure, T., Gjini, E., Stevens, J., Lane, W. J., … Wu, C. J. (2017). An immunogenic personal neoantigen vaccine for patients with melanoma. Nature, 547(7662), 217–221. 10.1038/nature22991

Piertney, S. B., & Oliver, M. K. (2006). The evolutionary ecology of the major histocompatibility complex. Heredity, 96(1), 7–21. 10.1038/sj.hdy.6800724

Pyke, R. M., Mellacheruvu, D., Dea, S., Abbott, C. W., Zhang, S. V., Phillips, N. A., Harris, J., Bartha, G., Desai, S., McClory, R., West, J., Snyder, M. P., Chen, R., & Boyle, S. M. (2021). Precision Neoantigen Discovery Using Large-scale Immunopeptidomes and Composite Modeling of MHC Peptide Presentation. Molecular & Cellular Proteomics: MCP, 20, 100111. 10.1016/j.mcpro.2021.100111

Racle, J., Guillaume, P., Schmidt, J., Michaux, J., Larabi, A., Lau, K., Perez, M. A. S., Croce, G., Genolet, R., Coukos, G., Zoete, V., Pojer, F., Bassani-Sternberg, M., Harari, A., & Gfeller, D. (2023). Machine learning predictions of MHC-II specificities reveal alternative binding mode of class II epitopes. *Immunity*, S1074761323001292. 10.1016/j.immuni.2023.03.009

Racle, J., Michaux, J., Rockinger, G. A., Arnaud, M., Bobisse, S., Chong, C., Guillaume, P., Coukos, G., Harari, A., Jandus, C., Bassani-Sternberg, M., & Gfeller, D. (2019). Robust prediction of HLA class II epitopes by deep motif deconvolution of immunopeptidomes. Nature Biotechnology, 37(11), 1283–1286. 10.1038/s41587-019-0289-6

Reynisson, B., Alvarez, B., Paul, S., Peters, B., & Nielsen, M. (2020). NetMHCpan-4.1 and NetMHCIIpan-4.0: Improved predictions of MHC antigen presentation by concurrent motif deconvolution and integration of MS MHC eluted ligand data. Nucleic Acids Research, 48(W1), W449–W454. 10.1093/nar/gkaa379

Sarkizova, S., Klaeger, S., Le, P. M., Li, L. W., Oliveira, G., Keshishian, H., Hartigan, C. R., Zhang, W., Braun, D. A., Ligon, K. L., Bachireddy, P., Zervantonakis, I. K., Rosenbluth, J. M., Ouspenskaia, T., Law, T., Justesen, S., Stevens, J., Lane, W. J., Eisenhaure, T., … Keskin, D. B. (2020). A large peptidome dataset improves HLA class I epitope prediction across most of the human population. Nature Biotechnology, 38(2), 199–209. 10.1038/s41587-019-0322-9

Sommer, S. (2005). The importance of immune gene variability (MHC) in evolutionary ecology and conservation. Frontiers in Zoology, 2(1), 16. 10.1186/1742-9994-2-16

Tadros, D. M., Eggenschwiler, S., Racle, J., & Gfeller, D. (2023). The MHC Motif Atlas: A database of MHC binding specificities and ligands. Nucleic Acids Research, 51(D1), D428–D437. 10.1093/nar/gkac965

Tareen, A., & Kinney, J. B. (2020). Logomaker: Beautiful sequence logos in Python. Bioinformatics, 36(7), 2272–2274. 10.1093/bioinformatics/btz921

The UniProt Consortium, Bateman, A., Martin, M.-J., Orchard, S., Magrane, M., Agivetova, R., Ahmad, S., Alpi, E., Bowler-Barnett, E. H., Britto, R., Bursteinas, B., Bye-A-Jee, H., Coetzee, R., Cukura, A., Da Silva, A., Denny, P., Dogan, T., Ebenezer, T., Fan, J., … Teodoro, D. (2021). UniProt: The universal protein knowledgebase in 2021. Nucleic Acids Research, 49(D1), D480–D489. 10.1093/nar/gkaa1100

Thibault, P., & Perreault, C. (2022). Immunopeptidomics: Reading the Immune Signal That Defines Self From Nonself. Molecular & Cellular Proteomics, 21(6), 100234. 10.1016/j.mcpro.2022.100234

Tran, E., Turcotte, S., Gros, A., Robbins, P. F., Lu, Y.-C., Dudley, M. E., Wunderlich, J. R., Somerville, R. P., Hogan, K., Hinrichs, C. S., Parkhurst, M. R., Yang, J. C., & Rosenberg, S. A. (2014). Cancer Immunotherapy Based on Mutation-Specific CD4+ T Cells in a Patient with Epithelial Cancer. Science, 344(6184), 641–645. 10.1126/science.1251102

Trolle, T., McMurtrey, C. P., Sidney, J., Bardet, W., Osborn, S. C., Kaever, T., Sette, A., Hildebrand, W. H., Nielsen, M., & Peters, B. (2016). The Length Distribution of Class I–Restricted T Cell Epitopes Is Determined by Both Peptide Supply and MHC Allele–Specific Binding Preference. The Journal of Immunology, 196(4), 1480–1487. 10.4049/jimmunol.1501721

Vita, R., Mahajan, S., Overton, J. A., Dhanda, S. K., Martini, S., Cantrell, J. R., Wheeler, D. K., Sette, A., & Peters, B. (2019). The Immune Epitope Database (IEDB): 2018 update. Nucleic Acids Research, 47(D1), D339–D343. 10.1093/nar/gky1006

Vyas, J. M., Van Der Veen, A. G., & Ploegh, H. L. (2008). The known unknowns of antigen processing and presentation. Nature Reviews Immunology, 8(8), 607–618. 10.1038/nri2368

Wagih, O. (2017). ggseqlogo: A versatile R package for drawing sequence logos. *Bioinformatics (Oxford*, England), 33(22), 3645–3647. 10.1093/bioinformatics/btx469

Ye, Y., Wang, J., Xu, Y., Wang, Y., Pan, Y., Song, Q., Liu, X., & Wan, J. (2021). MATHLA: A robust framework for HLA-peptide binding prediction integrating bidirectional LSTM and multiple head attention mechanism. BMC Bioinformatics, 22(1), 7. 10.1186/s12859-020-03946-z

Zeng, H., & Gifford, D. K. (2019). DeepLigand: Accurate prediction of MHC class I ligands using peptide embedding. Bioinformatics, 35(14), i278–i283. 10.1093/bioinformatics/btz330

